# Tyrosine kinase inhibitor independent gene expression signature in CML offers new targets for LSPC eradication therapy

**DOI:** 10.1101/2022.06.30.498287

**Authors:** Eduardo Gómez-Castañeda, Simon Rogers, Chinmay Munje, Joana Bittencourt-Silvestre, Mhairi Copland, David Vetrie, Tessa Holyoake, Lisa Hopcroft, Heather Jørgensen

## Abstract

2

Tyrosine kinase inhibitors (TKI) have revolutionised the treatment of CML. However, TKI do not eliminate the leukaemia stem cells (LSC), which can reinitiate the disease. Thus, finding new therapeutic targets in CML LSC is key to finding a curative treatment. Using microarray datasets, we defined a list of 227 genes which were differentially expressed in CML LSC compared to healthy controls but were not affected by TKI *in vitro*. Two of them, *CD33* and *PPIF*, are targeted by gemtuzumab-ozogamicin and cyclosporin A, respectively. We treated CML and control CD34^+^ cells with either drug with or without imatinib to investigate the therapeutic potential of the TKI-independent gene expression programme. Cyclosporine A in combination with imatinib reduced the number of CML CFC compared with non-CML controls, but only at supra-therapeutic concentrations. Gemtuzumab-ozogamicin showed a EC_50_ of 146ng/mL, below the plasma peak concentration of 630ng/mL observed in AML patients and below the EC_50_ of 3247ng/mL observed in non-CML cells. Interestingly, gemtuzumab-ozogamicin seems to promote cell cycle progression in CML CD34^+^ cells and demonstrated activation of the RUNX1 pathway in a RNAseq experiment. This suggests that targeting the TKI-independent genes in CML LSC could be exploited for the development of new therapies in CML.

**Simple summary:** Chronic myeloid leukaemia (CML) is initiated by a group of cancer cells called leukaemia stem cells (LSC). These LSC can survive current tyrosine kinase inhibitor (TKI) treatments and upon treatment withdrawal, are able to re-initiate the disease. Thus, eradicating the LSC would likely cure CML. In this study, we have identified a number of genes which expression is different between LSC and their healthy counterparts (haematopoietic stem cells) but are not affected by TKI treatment. We hypothesised that these genes may be potential therapeutic targets against LSC and used two different drugs, gemtuzumab-ozogamicin and cyclosporine A, to treat CML *in vitro*. We found that both drugs have a stronger effect on CML cells than in healthy cells. Therefore, we propose that the list of genes we identified could represent a novel source of therapeutic targets against CML.

## 4 Introduction

The introduction of tyrosine kinase inhibitors (TKIs) into clinical practice has greatly improved survival prognosis of patients diagnosed with chronic myeloid leukaemia (CML) [1]. However, TKIs are unable to eradicate the disease as leukaemic stem cells (LSCs) persist despite treatment [2-4]. This means that most patients need life-long therapy with its associated side effects[5], which is not only a financial and psychological burden, but upon which risk of resistance, disease recrudescence and progression are contingent.

TKI treatment, although effective at eradicating CML cells, has been shown to increase the proportion of quiescent LSCs [2-4,6], partially because of the potential for induction of quiescence and self-renewal gene expression [7,8]. Moreover, inhibition of BCR-ABL1 TK activity alone is not enough to eradicate CML progenitor cells [9] or LSCs [10]. Thus, it is possible that CML LSCs’ aberrant signalling is not entirely driven by BCR-ABL1 TK. For example, it has been reported that expression of *MIR10A*, a miRNA that is downregulated in CML cells, is not dependent on BCR-ABL1 TK but its downregulation promotes proliferation and cell growth in CML cells [11]. As TKI monotherapy is insufficient to eradicate CML LSCs, this suggests that several genes independent of BCR-ABL1 TK may be supporting these BCR-ABL1 TK independent cells.

This led to the hypothesis that there is a gene expression programme in CML LSCs independent of the BCR-ABL1 TK activity that is required for their survival. Previous reports have investigated the presence of deregulated genes in CML cells that are not corrected by TKI in therapy resistant cells. For example, the β-catenin protein levels of CML CD34^+^ cells from patients resistant to more than one TKI are unaffected by TKI treatment *in vitro* [12]. Additionally, MYC and p53 pathways are deregulated in CML LSC regardless of the responsiveness to TKI treatment and, upon investigation, were shown not to be targeted by TKI [13].

Based on the current evidence, we hypothesised that treatment naïve CML CD34^+^ cells possess a gene expression programme independent of the BCR-ABL1 TK which allows them to persist following TKI treatment and that the components of this programme could be targeted to eradicate the leukaemic clone. By investigating CML CD34^+^ cells treated with TKI and untreated, as well as CML LSC and normal haemopoietic stem cells (HSC), we identified a list of 227 genes that are TKI independent (TKI-independent) and potentially BCR-ABL1 TK independent. This list includes *CD33*, a myeloid cell surface marker, *PPIF*, a cell death regulator, and *ERG*, a transcriptional factor involved in HSC development and maintenance. We used gemtuzumab-ozogamicin (GO), an anti-CD33 conjugated monoclonal antibody approved by the Food and Drug Administration (FDA) and the European Medicines Agency (EMA) for the treatment of acute myeloid leukaemia [14] and cyclosporin A (CsA), a blocker of PPIF activity [15] for targeting CML CD34^+^ cells under physiological conditions *in vitro* (Figure 1a). *ERG* downregulation mimics *MYC* overexpression and it has been shown that bromodomain and extraterminal (BET) inhibitors restore normal expression in ERG deficient mice [16]. BETi have already been successfully shown to eliminate CML LSC in our lab [13].

**Figure 1.**
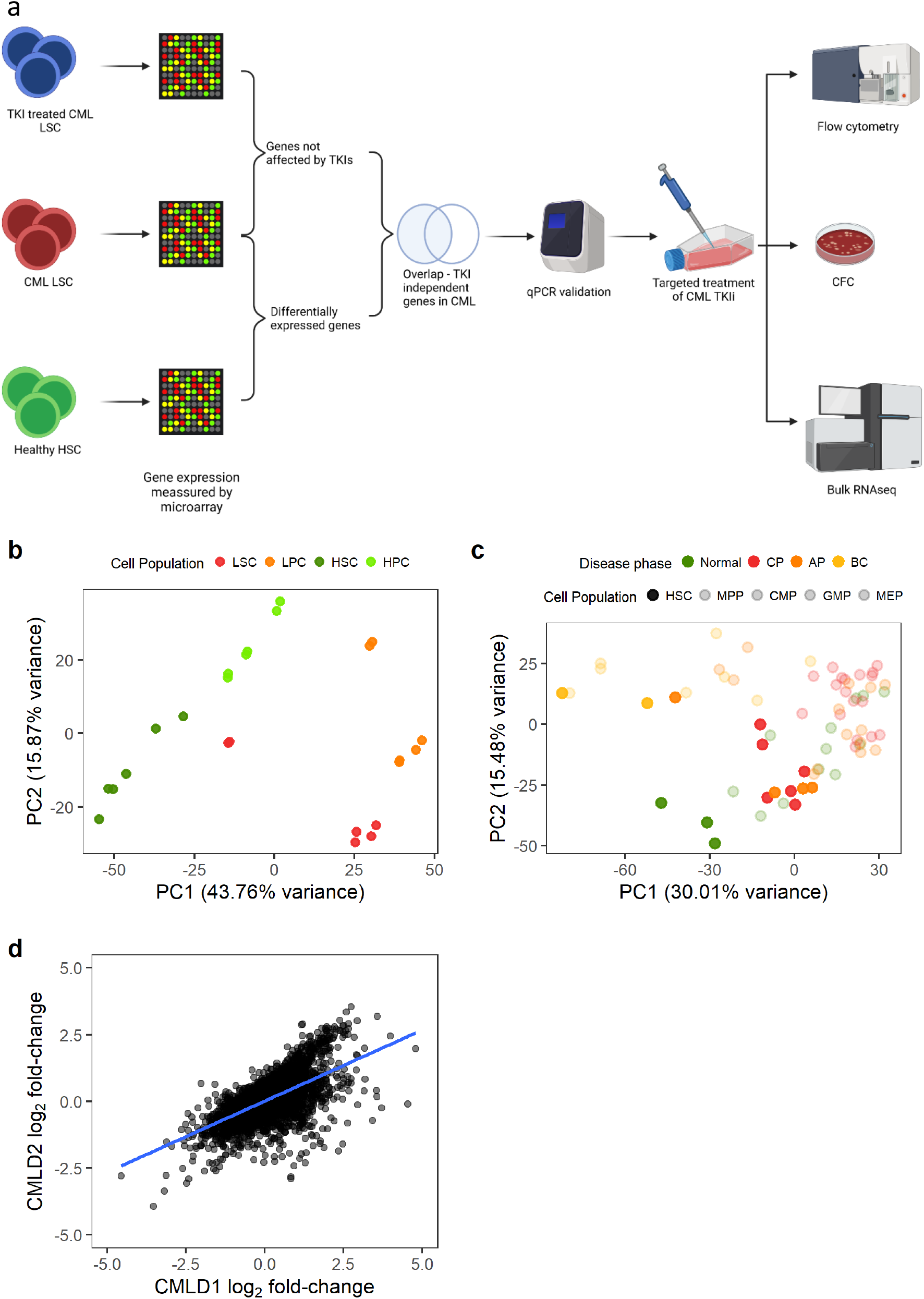
Gene expression changes in CML LSC are consistent across datasets. (a) Experimental design. We identified genes differentially expressed in CML LSC but not affected by TKI treatment (expression measured by microarrays) and, selected two targets for targeted therapy against CML LSC in combination with TKI. The effect of such therapies was studied by flow cytometry (cell cycle), CFC and RNAseq. (b) PCA plot of CMLD1. Each dot represents a microarray (including technical duplicates). Technical duplicates were close to each other. The samples clustered following two axes: CML/normal and stem/progenitor cells. (c) PCA plot of CMLD2. Each dot represents one microarray (no technical replicates in this dataset). CML-CP LSC co-localise with normal progenitor cells, suggesting a progenitor-like phenotype in CML-CP LSC. (d) Gene expression log_2_ fold-changes between CML LSC and normal HSC were significantly correlated between CMLD1 and CMLD2 (Pearson’s R = 0.63). Each dot represents an individual gene.

## 5 Material and methods

### 5.1 Code availability

The code used for the data analysis can be found in GitHub <u>EduardoGCCM/Gomez-Castaneda2022_TKI-independent</u>.

### 5.2 Primary patient material

All samples were collected after written informed consent from the patients. The project had ethical approval from the West of Scotland Research Ethics Committee 4 (REC reference: 15-WS-0077). Cells were processed for cryopreservation from peripheral blood or leukapheresis from patients diagnosed with CML, and other haematological malignancies or healthy donors (‘non-CML’ including allogeneic haematopoietic stem cell donors). Patients’ age, biological sex and, response to imatinib (IM) are summarised in Tables S1 (CML) and S2 (non-CML). Cells were enriched for CD34^+^ using magnetic activated cell sorting (MACS) or CliniMACS.

## 5.3 Microarray analysis

Data analysis was performed using R 4.1.1 running under MS Windows 10 21H1 unless otherwise stated.

Microarray intensities were normalised using the RMA algorithm [17-19] implemented in the *oligo* package [20]. Microarray chip annotations were summarised at the transcript level using the *getNetAffx* function. Microarray differential gene expression was calculated using *limma* [21,22]. Statistically significant probe sets were annotated to gene names (HGNC symbols) with *biomaRt* [23] connecting to the Ensembl release August 2020 [24].

The microarray datasets used for comparison of CML and non-CML stem cells (LSC/HSC) were CMLD1 (Array Express accession number E-MTAB-2581) [13] and CMLD2 (GEO accession number GSE47927) [25]. CMLD1 comprises of LSC/HSC (CD34^+^CD38^-^) and progenitor cells (CD34^+^CD38^+^) from three CML patients and three non-CML patients. Samples from CMLD1 were run in technical duplicates. Only the LSC/HSC were used for the differential expression analysis and technical replicates were accounted using the *duplicateCorrelation* function from *limma*. CMLD2 comprises of a collection of cell populations of varying maturity — LSC/HSC (CD34^+^CD38^-^CD90^+^), MPP (CD34^+^CD38^+^CD90^-^), CMP (CD34^+^CD38^+^CD123^+^CD45RA^+^), GMP (CD34^+^CD38^+^CD123^+^CD45RA^low^) and MEP (CD34^+^CD38^+^CD123^-^CD45RA^-^) — sorted from healthy donors (n=3), chronic phase (n=6), accelerated phase (n=4) and myeloid blast crisis CML patients (n=2). Only LSC/HSC from healthy and CML chronic phase were used for the analysis. A q-value smaller than 0.1 (Benjamini-Hochberg correction [26]) was considered statistically significant.

Comparison between TKI treated and untreated cells was made by comparing the samples in TKID1 (Array Express accession number E-MTAB-2594) [27]. Gene expression was measured at baseline (0h) and after eight hours of treatment (8h) for CD34^+^CD38^-^ and after 7 days treatment (7d) for CD34^+^ cells. The cells were treated with the clinically achievable drug concentrations of 5μM imatinib (IM), 150nM dasatinib or 5μM nilotinib (one replicate each); a second dose of the drug at the same concentration was applied during the 4^th^ day of treatment. The cells were sorted again for live cells after 7d of treatment before RNA extraction. Statistical significance for no change was assessed using an equivalence test [28].

All the microarrays analysed were performed using Affymetrix HuGe 1.0 ST chips and RNA was extracted using RNeasy Micro Kit (Qiagen) when the number of isolated cells was less than 5×10^5^, and RNeasy Mini Kit (Qiagen) when the number of isolated cells was between 5×10^5^ and 1×10^7^.

### 5.4 RNA sequencing

RNA was extracted using the Arcturus PicoPure kit for patients CML423 and CML460 and using the RNAeasy Micro kit for patient CML441. RNA was reverse transcribed using the SMART-Seq v4 Ultra Low Input RNA Kit for Sequencing (Takara, Saint-Germain-en-Laye) by Glasgow Polyomics. cDNA library preparation was performed using Nextera library preparation kit by Glasgow Polyomics. RNA sequencing was performed using an Illumina HiSeq 4000 sequencer. The dataset can be found in GEO (GSE198576).

### 5.5 Bulk RNA-seq analysis

RNAseq quality of sequencing was assessed using *FastQC* [29], and sequences were trimmed using *cutadapt* [30] with default parameters on pair-end mode removing the last base of each read, filtering out reads shorter than 20bp. The release v31 of the GRCh38.p12 human genome was used as reference and reads were aligned to the genome using *STAR [31]*. Count matrix was generated using *featureCounts* (Liao et al. 2013). Differential gene expression was calculated using *edgeR* [32]. A q-value smaller than 0.1 (Benjamini-Hochberg correction [26]) was considered statistically significant.

### 5.6 Validation of the TKI-independent signature

Cells were seeded in serum free media (SFM) in a 6-well plate and grouped as no drug control (NDC) or IM treated, as previously described for the TKID dataset [27]. IM was added at a final concentration of 5μM. Cells were cultured at 37°C and 5% CO2 for 7 days. IM was added again at day 4 without washing the cells. On day 7 cells were sorted for viable cells (DAPI^-^) by flow cytometry using a BD FACS Aria with Diva software and used for RNA extraction. RNA was extracted using RNeasy Micro Kit (Qiagen) when the number of isolated cells was less than 5×10^5^, and RNeasy Mini Kit (Qiagen) when the number of isolated cells was between 5×10^5^ and 1×10^7^.

### 5.7 qPCR analysis

qPCR was performed using a Fluidigm 48.48 PCR chip following the manufacturers protocol.

The cDNA of each sample was pre-amplified for 18 cycles using the PCR multiplex PCR kit (Qiagen). Each reaction contained a pool of all the primers of interest at 50nM each and a maximum 12.5ng of cDNA (some samples yielded very low concentrations of RNA and higher concentrations were not possible). The polymerase was activated at 95ºC for 15 minutes and each cycle comprised of 30 seconds of denaturation at 94ºC, 90 seconds of annealing at 60ºC and 60 seconds of extension at 72ºC. A final extension of 30 minutes at 72ºC was performed. The samples were treated with 0.5U/μL of exonuclease I (New England Biolabs, Ipswich, MA, USA) for 30 minutes at 37ºC. The enzyme was inactivated at 80ºC for 15 minutes. The samples were diluted 1:5 and stored.

The 48.48 chip was primed with control line fluid in the IFC controller MX and each of the primer wells was filled with a 5μL solution containing 1X assay loading reagent, DNA suspension buffer and 5μM of each of the primers of the pair assigned to the well. Each sample was loaded with 5μL of 1X SsoFast™ EvaGreen Supermix with low ROX (Bio-Rad), 1X DNA binding dye sample loading reagent (Fluidigm) and 45% v/v of the pre-amplified cDNA assigned to the well. The reaction in the Biomark activated the enzyme at 95ºC for 1 minute and performed 30 cycles of denaturation at 96ºC for 5 seconds and annealing and extension at 60ºC for 20 seconds. A melting curve was generated at the end of the qPCR for every reaction.

Relative expression of the test genes was calculated by subtracting the mean of the Ct values of the reference genes (*ENOX2, GAPDH, RNF20*, and *TYW1*) to the Ct value of the test gene within each sample (ΔCt). These ΔCt values were used as normalised gene expression values and differential gene expression was calculated using *limma* (Ritchie et al., 2015). Genes were considered to be differentially expressed when the BH-adjusted p-value was lower than 0.1. The confidence interval of the ΔΔCt (log2 fold change) was also calculated by *limma*. A gene was considered non-changing when its median ΔΔCt was within the interval -0.5 to 0.5.

### 5.8 Drug response experiments

CD34^+^ primary cells were resuspended at a density of 2×10^5^cells/mL in SFM + physiological growth factors (0.2ng/mL SCF, 1ng/mL G-CSF, 0.2ng/mL GM-CSF, 1ng/mL IL6, 0.05ng/mL LIF, 0.2ng/mL MIP1α) in the presence or absence of 2μM IM and/or different concentrations of either GO (10, 30, 100, 300 and 1000ng/mL) or CsA (0.3, 1, 3, 5, 10 and 30 μM) and cultured at 37°C and 5% CO_2_. Treatment was delivered in three different regimens:

- 72 hours of GO or CsA with or without IM.
- 72 hours IM (or no drug) followed by 72 hours of either GO or CsA.
- 72 hours of either GO or CsA followed by 72 hours of IM (or no drug).

Following 72h the cells were washed in PBS and centrifuged for 10 minutes at 300g three times in order to washout the first drug. After treatment, viable cells were manually counted using trypan blue dye exclusion using a haemocytometer and used for downstream experiments.

### 5.9 Colony forming cell assays

For colony forming cell (CFC) assay, 3,000 cells were mixed with 3mL of Methocult^®^ H4034 (Stem Cell Technologies). This mix was then split evenly between two 35mm^2^ plates covering the entire surface of the plates. All the plates from each sample were placed inside a 23.5cm^2^ plate and 2 plates containing just water were added to avoid the Methocult drying. Cells were cultured for no less than 9 days at 37°C and 5% CO_2_ before analysis.

### 5.10 Cell cycle analysis

Cells were fixed using 80% ethanol at -20°C and stored at 4°C before analysis. DNA was stained using DRAQ7 dye and analysed on a BD FACS Aria (BD Biosciences) with Diva software. Data analysis was performed with FlowJo 10.7.2.

### 5.11 Other data analysis

Unless otherwise stated data analysis was performed using R 4.1.1 on Windows 10 21H1. A q-value smaller than 0.1 was considered statistically significant (Benjamini-Hochberg correction [26]. Drug response curves were calculated using the *drc* package [33].

## 6 Results

### 6.1 CML LSC possess a TKI-independent gene expression signature

We first identified the genes that were differentially expressed (DE) between CML LSC and non-CML HSC (Figure 1a). We used two previously generated microarray datasets comparing non-CML to treatment naïve CML cells. As described in the methods section, CMLD1 [13] and CMLD2 [25] were generated using the same Affymetrix chips and the most primitive populations (HSC/LSC) were sorted for CD34^+^CD38^-^ and CD34^+^CD38^-^CD90^+^, respectively. The first 2 principal components (PC) of CMLD1 showed a clear separation between CML and non-CML, as well as between stem (CD34^+^CD38^-^) and progenitor cells (CD34^+^CD38^+^), as expected (Figure 1b). The first two PC of CMLD2 showed non-CML HSC in a distinct position, while chronic phase (CP) CML and most accelerated phase (AP) CML LSC clustered with non-CML progenitors, suggesting that CP and AP CML LSC have a progenitor-like molecular phenotype, as previously reported [34]. The blast crisis (BC) LSC and one AP LSC sample clustered together, as previously shown using scRNAseq [6]. When comparing CML LSC against non-CML HSC in CMLD1 we identified 4,505 DE genes, while the same comparison in CMLD2 identified 2,344 DE genes. In order to investigate only genes with a high confidence of being DE in CML LSC compared with non-CML HSC, we used the 1,497 genes that were DE in both datasets. This overlap (or higher) was very unlikely to happen by chance (p<0.001, hypergeometric distribution). Additionally, we investigated if the global changes in the transcriptome were conserved across datasets (Figure 1d). We found significant positive correlation in both cases (p<0.001; R_1d_=0.63; Pearson’s correlation).

Then, we identified the genes not affected by TKI treatment using the TKID dataset. TKID was generated with the same Affymetrix chips as CMLD1 and CMLD2 and contains gene expression data of treatment naïve chronic phase CML samples at baseline, (CD34^+^CD38^-^ cells) and after 7 days *in vitro* TKI treatment (7d, CD34^+^ cells). The PCA suggests a strong patient effect (patients indicated by colour), with the cells treated with TKI for 7d being different from the cells at baseline or the cells after early exposure (Figure 2a). As we were trying to identify genes not affected by TKI treatment (as a proxy for genes independent from BCR-ABL1 TK), we decided to assess TKI independent (TKI-independent) genes using an equivalence test, as previously described [28]. As equivalence tests require a threshold or margin of equivalence to be set by the research team, we decided to use the expected technical variation. Thus, we calculated the log_2_ fold-changes of technical replicates present in CMLD1 and TKID in order to identify the expected differences between biologically identical samples. As a p-value threshold of 0.05 is commonly used as a reference for significant change, we selected the percentiles 2.5 and 97.5, as the threshold for change. These percentiles were -0.486 and 0.487 log_2_ fold-change, which we rounded to -0.5 and 0.5 for the remaining calculations. Thus, any log_2_ fold-change whose 95% confidence interval crossed zero and was contained within -0.5 to 0.5 was considered non-changing. We identified 4,913 non-changing genes after 7d of TKI treatment *in vitro*.

**Figure 2.**
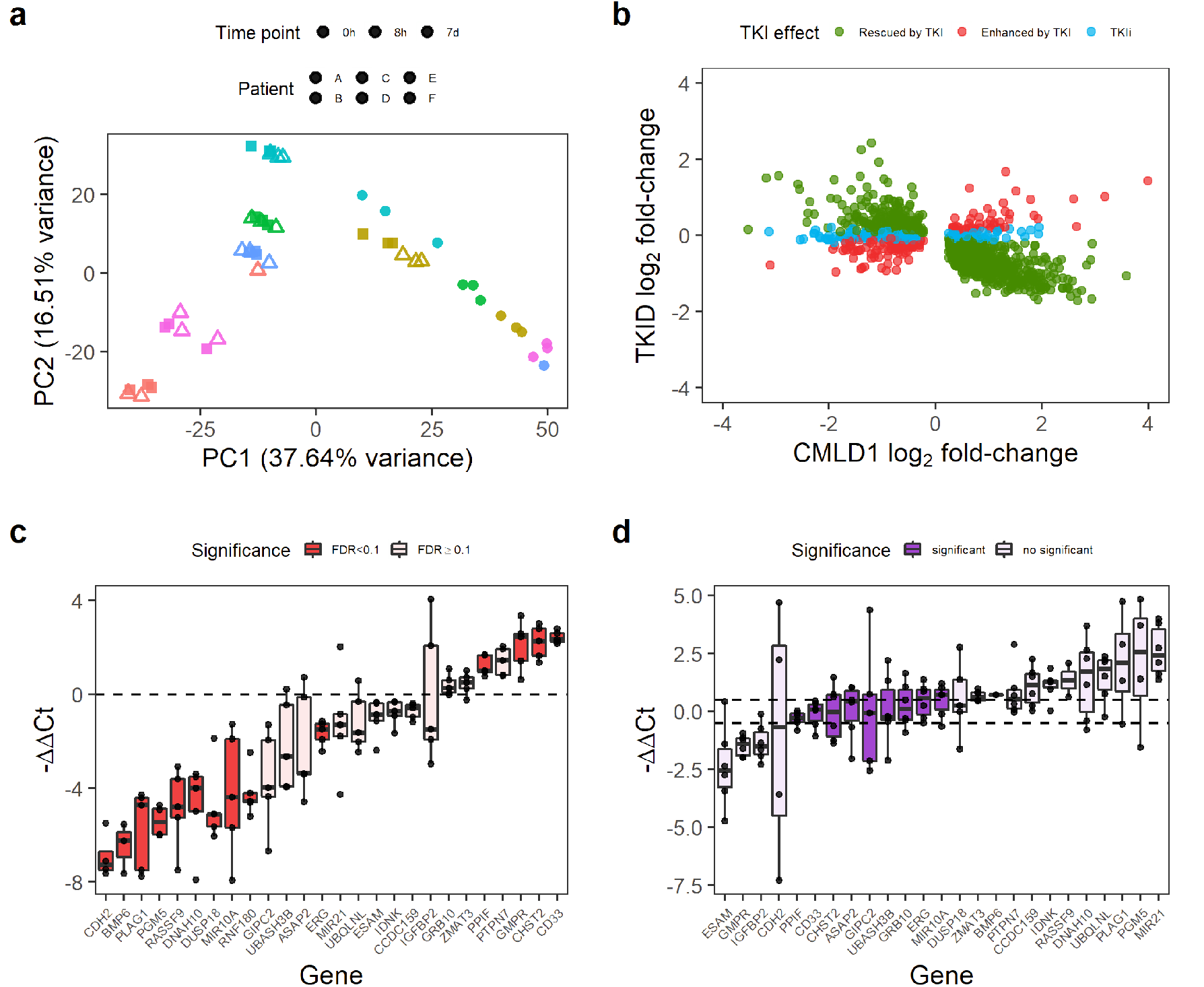
Some genes differentially expressed in CML LSC are not affected by TKI treatment. (a) PCA of the TKID dataset. Interpatient variability and treatment (7d vs 0h/8h) seem to have a major contribution to the variance of the dataset. Each point represents one microarray. (b) Correlation of gene expression log2 fold-changes between TKID and CMLD1. Although most genes show a negative correlation after 7d of TKI treatment compared with CML vs Normal (normal expression “rescued by TKI”), a minority of genes were not affected by TKI treatment (TKI independent or TKIi) and some genes were further deregulated into the same direction than in CML vs Normal (“enhanced by TKI”). (c) Gene expression differences between CML-CP CD34^+^ cells and normal CD34^+^ cells as measured by qPCR. Each dot represents the relative expression of one CML sample compared with the mean expression of the normal samples for each gene. Differential gene expression was calculated with limma and an FDR<0.1 was considered significant. (d) Gene expression differences between CML-CP CD34^+^ cells treated with 5μM IM (7d) and NDC (7d). Relative gene expression was calculated per patient (paired-analysis) and a mean log2 fold-change (or ΔΔCt) between -0.5 and 0.5 was considered significant no change. Each dot represents a patient.

74.2% (1,111/1,497) of the consistently DE genes between CML and non-CML were rescued by TKI (i.e., the effect of TKI treatment was opposite to that of CML transformation). However, the expression of 159 genes was further de-regulated after TKI treatment (e.g., genes already upregulated in CML were further upregulated) and 227 genes were not affected by TKI treatment (Figure 2b). We termed these latter genes TKI-independent genes and focused on them as potential therapeutic targets for combination treatments with TKIs. The 227 TKI-independent genes can be found in Table S3.

We selected a group of 26 genes, which had an absolute log_2_ fold-change of at least 0.7 in CMLD1 (only *CHST2* and *IDNK* had smaller fold-changes) and had biological relevance, for validation by qPCR in an independent cohort of five CML-CP patients and two healthy donors. The qPCR experiment confirmed that of the 26 genes, 14 were DE between CML and non-CML (Figure 2c) and 9 were not affected by IM (Figure 2d). *PPIF, ERG, MIR10A, CD33* and *CHST2* were TKI-independent in the validation experiment. As PPIF and CD33 can be targeted by existing drugs (CsA and GO, respectively), we focused on these two genes for targeting the TKI-independent signature in CML LSCs.

### 6.2 Anti-CD33 therapeutic antibody can be used to specifically eradicate CML CD34^+^ cells

GO is a conjugated monoclonal antibody that releases the toxic antibiotic calicheamicin intracellularly after it binds its target, CD33. Calicheamicin produces double strand breaks in the DNA, leading to cell death. Currently, GO is approved for the treatment of AML by both the FDA and the EMA [35]. Thus, we consider it a potential candidate for re-purposing. Furthermore, a previous study found GO to be effective at eradicating CML mononuclear cells [36]. We also confirmed by flow cytometry that expression of CD33 on the cell surface did not significantly change after treating the cells with IM (Figure S1).

We treated CML and non-CML CD34^+^ cells using three different regimens *in vitro*, to mimic potential clinical scenarios when used in combination with IM: GO or GO+2μM IM for 72h treatment (GO_72_); 72h IM or NDC followed by 72h GO treatment (GO_IMGO_); or, 72h GO followed by 72h IM or NDC (GO_GOIM_).

Using the GO_72_ regimen, CML CD34^+^ cells had an EC_50_ of 146ng/mL when treated with GO alone but 195ng/mL when combined with IM (Figure 3a). These EC_50_s were not statistically significantly different from each other and thus the effect of GO and IM can be considered independent. These concentrations are achievable *in vivo* with the current recommended dosing for the treatment of AML (plasma C_max_ of 630ng/mL) [35].

**Figure 3.**
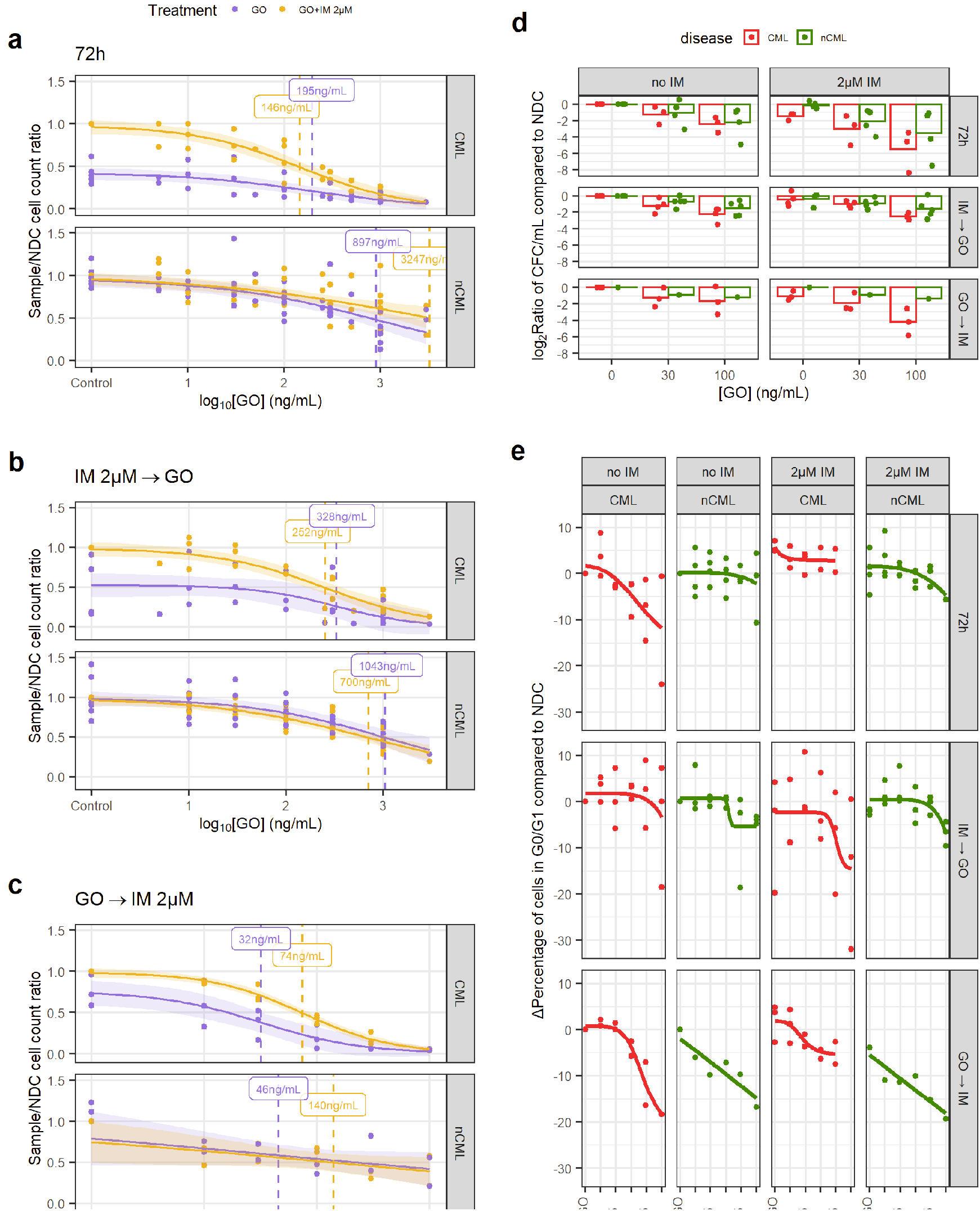
GO is effective at eradicating CML CD34^+^ cells. (a) Dose-response curves for the GO_72_ treatment regimen. (b) Dose-response curve for the GO_IMGO_ treatment regimen. (c) Dose-response curve for the GO_GOIM_ treatment regimen. (d) The reduction in the number of CFC was calculated dividing the total number of CFC in the original liquid culture after treatment and dividing this number by the number of CFC in the original liquid culture of the corresponding NDC (patient matched). These ratios were log_2_-transformed. A linear model fitted using GO_72_ and GO_IMGO_ data showed that GO significantly reduces CFC numbers in CML samples more than in nCML samples. (e) Change in the percentage of quiescent (G_0_/G_1_) cells after treatment at different concentrations and regimens of GO. Cell cycle was assessed using DRAQ7 DNA staining (flow cytometry). (All) The shadowed area represents the 95% confidence interval and the dashed line the EC_50_ (value given in the figure). Each dot represents an individual patient sample (across different concentrations). Curves and EC_50_ were calculated using R’s *drc* package.

Additionally, GO on non-CML controls had an EC_50_>3,000ng/mL that decreased to 897ng/mL in combination with IM (Figure 3a), both being significantly higher than respective EC_50_ for CML cells. These results show that GO has a wide therapeutic window and that non-CML EC_50_ remains higher than clinically achievable plasma peak concentrations in patients under the current dosing recommendation for GO for AML.

The GO_IMGO_ regimen showed a similar trend, with both GO alone and in combination with IM having significantly lower EC_50_ in CML than in non-CML and within *in vivo* achievable concentrations (Figure 3b). EC_50_ in non-CML remained higher than peak plasma concentrations in patients.

The GO_GOIM_ regimen showed an overall reduction in EC_50_ for all conditions. Although the number of samples tested with this regimen is lower than the other two, it might suggest that GO cytotoxic effect may persist after the drug has been removed and washed out from the culture.

In order to investigate the effect of GO on stem and progenitor cells (SPC), CFC counts and cell cycle analysis were performed. When comparing the treated/NDC log_2_ CFC ratios, CML cells consistently showed a greater decrease of CFCs than non-CML in response to drug, which was significant in a linear model (1.46-fold bigger decrease than non-CML, p=0.024; Figure 3d). GO_GOIM_ was not included in the linear model due to the imbalance of CML/non-CML sample number. This suggests that CML SPC are more susceptible to GO than non-CML SPC.

IM is known for inducing quiescence in SPC, protecting CML LSC from eradication. Thus, combining IM with a drug that pushes CML LSC towards cell cycle may increase their vulnerability to IM. Using DNA staining, we observed a decrease in the number of CML cells in either G_0_ or G_1_ when treated with either GO_72_ or GO_GOIM_ regimens, but not with GO_IMGO_. IM treated CML cells showed an increase in cells in G_0_/G_1_ phases that was only modestly decreased after GO treatment (Figure 3e). Under the GO_IMGO_ regimen this was not observed, but it could be due to the increased cell concentration in the culture after 72h without treatment, which could affect either the growth potential of the cells or reduce the drug to target ratio, or other culture artifacts. The effect of GO on cell cycle stage of non-CML cells was minimal, especially at concentrations equal to, or lower than, CML EC_50_.

### 6.3 CsA targets CML SPC when in combination with IM

CsA is an immunosuppressant drug that has been used for the treatment of transplant rejection and graft versus host disease (GvHD) since the 1980s. It is known to inhibit the activation of cytotoxic T-cells. However, it has several side effects due to its broad mechanism of action [37]. Interestingly, CsA has also been proposed as a treatment to prevent necrosis in ischemic events due to its ability to inhibit the activity of PPIF (also known as cyclophilin D) and its interaction with p53 [15]. Thus, we tested whether inhibiting PPIF, which is upregulated in CML LSC compared to non-CML HSC, would have any benefit in the treatment of CML.

Similarly to the experiments using GO, we treated CML and non-CML CD34^+^ cells using three different regimens *in vitro*, to mimic potential clinical scenarios when used in combination with IM: CsA or CsA+2μM IM 72h treatment (CsA_72_); 72h IM or NDC followed by 72h CsA treatment (CsA_IMCA_); or, 72h CsA followed by 72h IM or NDC (CsA_CAIM_).

In all three regimens, the EC_50_ of CsA, either alone or in combination with IM was similar, suggesting an effect independent of IM. However, CsA’s EC_50_ for CML and non-CML were similar at 3,000-5,000nM and over the 1,000nM peak concentration usually achieved in the clinical practice (Figures 4a-c) [38].

**Figure 4.**
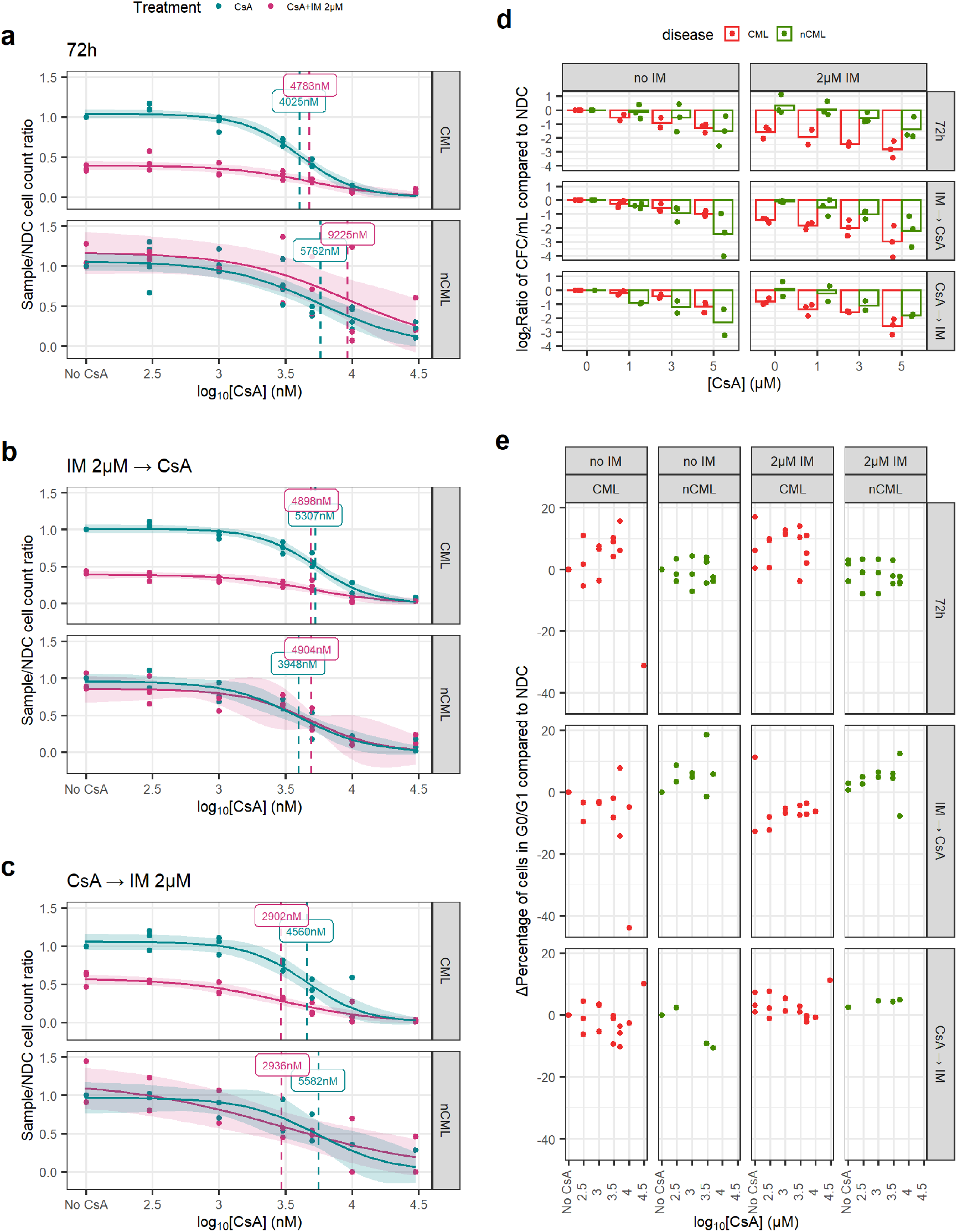
CsA eradicates CML CFC in combination with IM. (a) Dose-response curves for the CsA_72_ treatment regimen. (b) Dose-response curve for the CsA_IMCA_ treatment regimen. (c) Dose-response curve for the CsA_CAIM_ treatment regimen. (d) The reduction in the number of CFC was calculated dividing the total number of CFC in the original liquid culture after treatment and dividing this number by the number of CFC in the original liquid culture of the corresponding NDC (patient matched). These ratios were log_2_-transformed. Using all treatment regimens, a linear model showed that CsA significantly decreases more the number of nCML CFC than CML CFC in the absence of IM. However, in combination with IM CsA significantly reduces more the number of CFC in CML than in nCML. (e) Change in the percentage of quiescent (G_0_/G_1_) cells after treatment at different concentrations and regimens of CsA. Cell cycle was assessed using DRAQ7 DNA staining (flow cytometry). (All) The shadowed area represents the 95% confidence interval and the dashed line the EC_50_ (value given in the figure). Each dot represents an individual patient sample (across different concentrations). Curves and EC_50_ were calculated using R’s *drc* package.

Similarly, CsA alone has a greater effect on the number of non-CML CFCs than on CML (p=0.017). However, this trend is reversed when the cells have also been treated with IM, independently of the treatment regimen (p<0.001) (Figure 4d). This potentially means that CsA synergies with IM when targeting CML SPC.

In contrast with GO, CsA did not induce cell cycle progression and regimens/conditions showed increased cell cycle arrest with increasing concentrations of CsA, with or without IM (Figure 4e).

### 6.4 Transcriptomic changes after GO, CsA and IM treatments

Our results suggest that GO, and to some extent CsA, may be effective in eradicating CML CD34^+^ cells. To investigate the mechanisms of action by which GO and CsA, as well as IM, eradicate CML we performed bulk RNAseq from cells treated with 100ng/mL GO, 100ng/mL GO + 2μM IM, 2μM IM and 3μM CsA for either GO_72_ or CsA_72_ treatment regimens. As expected, the GO+IM combination treatment had the biggest impact on gene expression compared to the other conditions (Figure 5a). To increase the sensitivity for gene expression changes in the other treatments, differential gene expression from the single-agent treatments was performed on data normalised in the absence of the GO+IM combination. Removing the GO+IM combination allowed us to observe different clusters of samples according to their single-agent treatment (Figure 5b). Interestingly, none of the single-agent clusters overlapped and the NDC is projected to the centre of the plot when using the first two PCs. This may suggest that all three single agents affect different transcriptional pathways.

**Figure 5.**
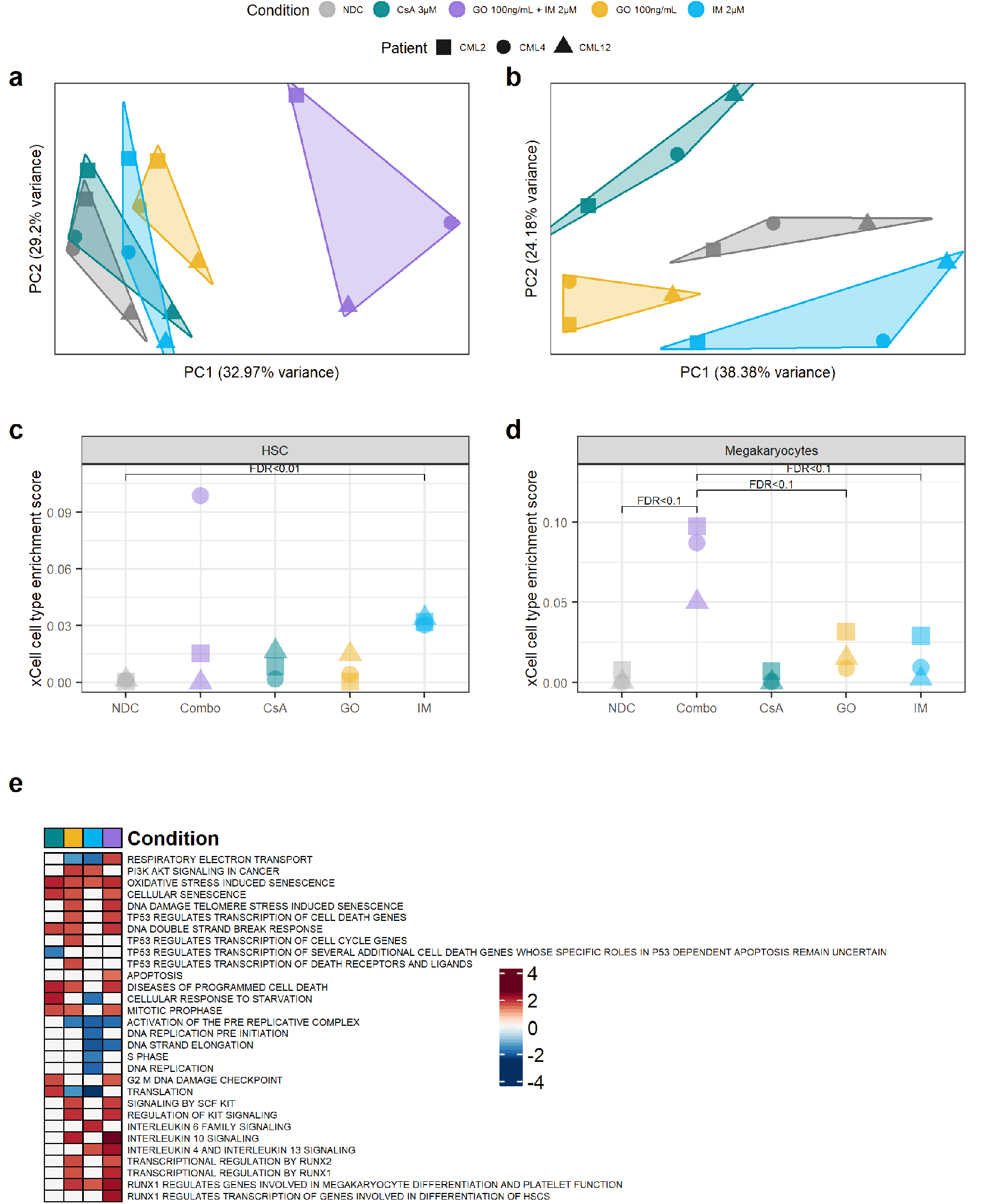
Targeting the TKIi signature alters the transcriptome of CML CD34^+^ cells. (a) PCA plot of all the samples sequenced. The transcriptomic changes caused by the GO+IM combination explain most of the variability of the dataset. (b) PCA plot after removing the GO+IM combination samples. This allows for a better understanding of the differences between the single treatments, which seem to be independent of each other (each treatment is projected in a different direction from the NDC). (c) Comparison of the HSC enrichment score generated by xCell for each sample. IM treated samples were significantly enriched by HSC compared to the NDC (FDR<0.01). (d) Comparison of the megakaryocyte enrichment score generated by xCell for each sample. The GO+IM was borderline significantly enriched on megakaryocytes compared either to the NDC or to GO/IM single treatment (FDR<0.1). (e) Shortlist of the Reactome pathways significantly enriched in any of the conditions by GSEA. White squares represent non-significant pathways (independently of the enrichment score). Blue squares represent downregulated pathways while red squares represents upregulated pathways. Full list of pathways can be found in Figure S3. (all) Each shape represents an individual patient while each colour represents an individual treatment.

Compared to the NDC, the GO+IM combination had 3,789 DE genes, GO single-agent 1,268 DE genes, CsA single-agent 695 DE genes, and IM single-agent 1,118 DE genes. Of those, the GO+IM combination had 57 DE TKI-independent genes, GO single-agent 19 DE TKI-independent genes, CsA single-agent 9 DE TKI-independent genes and, IM single-agent 17 DE TKI-independent genes (Table S4) (all DE genes available in Tables S5 to S8). The number of DE TKI-independent genes was the one expected by chance in all conditions, as reported by hypergeometric distribution (p_combination_=0.82, p_IM_=0.72, p_GO_=0.71, p_CsA_=0.15), suggesting that none of the treatments specifically targets all the TKI-independent genes. This is expected by GO and CsA, as they are targeted therapies against only one TKI-independent gene each, *CD33* and *PPIF*, respectively. However, we expected the TKI-independent genes to be under-represented in the IM treated cells.

Therefore, we next investigated why TKI-independent genes were not under-represented after IM treatment. This includes *CD33* and *ERG*, which were found to be TKI-independent genes also by qPCR. As *ERG* was upregulated after IM treatment we hypothesised that IM enriched for LSC in culture, as previously reported [7]. Thus, we used xCell [39] to calculate the HSC/LSC enrichment score in all treatment conditions and compared them to the NDC as well as single agent GO and IM to the combination treatment. xCell allows for the estimation of cell type enrichment across different samples using gene signatures derived from well annotated datasets, such as the Human Primary Cells Atlas [40]. Interestingly, IM treated cells showed a significant enrichment in HSC/LSC compared to the NDC (FDR=0.01) (Figure 5c, Table S9). As this enrichment was not detected in the TKID microarray dataset (Figure S3), the DE TKI-independent genes are likely DE due to the change in cell-type composition in the IM treated cells compared to the NDC, similar to previous reports [7].

Following this, we investigated if any other cell type expression signatures were enriched in any of the conditions. We found that the GO+IM combination was borderline significantly enriched in megakaryocytes when compared with the NDC or any of the two drugs alone (FDR<0.1) (Figure 5d, Table S10), suggesting that the combination treatment may promote HSC/LSC differentiation towards the megakaryocytic lineage. This difference was not detected in TKID (Figure S4).

Additionally, gene expression changes (compared to NDC) for each condition were analysed for pathway enrichment analysing using GSEA. We found that CsA upregulates pathways related to cell death while it downregulates pathways associated with p53 (Figure 5e). This was expected as the described mechanism of CsA is the inhibition of PPIF, which is known to interact with p53, promoting necrosis, and with anti-apoptotic BCL2, preventing programmed cell death [15]. IM also showed an expected effect, with cell cycle related pathways being downregulated after IM treatment. GO showed an enrichment of pathways involved in DNA double strand breaks response, which was expected because of the mechanism of action of calicheamicin. Interestingly, GO treated cells also showed an enrichment of pathways involved on cell differentiation through RUNX1/2 and KIT, especially in combination with IM (Figure 5e and Figure S2). The results from the GSEA support the effects of the drugs observed experimentally and the proposed mechanisms of action of GO and CsA.

## 7 Discussion

The existence of a BCR-ABL1 TK independent gene expression signature that allows CML LSC to persist after TKI treatment has been long hypothesised [2-4,9,10]. Indeed, different signalling pathways, such as β-catenin [12] MYC and p53 pathways [13] have been shown to be unaffected by TKI treatment, but to be deregulated in CML LSC compared with their healthy HSC counterparts. In this study we define a list of 227 genes (Table S3) that are differentially expressed in CML CD34^+^ cells but are not affected by TKI treatment. Despite the small sample size and the lack of *in vivo* post-TKI patient cells, this list suggests the existence of a transcriptomic signature in CML that is not affected by TKI and which is likely independent from the TK activity of BCR-ABL1 (although not necessarily BCR-ABL1 independent).

We have also shown that targeting the gene products from our list with existing compounds, such as GO and CsA, could open new avenues for the treatment of CML. GO is currently marketed for the treatment of AML and it seems to have low long-term toxicity on healthy HSCs at its recommended dose [35]. This low HSC toxicity is supported by our experiments as we observed an EC_50_>3,000ng/mL in non-CML CD34^+^ cells (897ng/mL in combination to 2μM IM). On the other hand, we have shown that GO is effective at reducing the number of CML CD34^+^ cells. Previous studies have shown that GO is effective against CML chronic phase mononuclear cells *in vitro* [36] and it has been used with partial success when treating patients with TKI resistant CML blast crisis [41-43]. Here we show that GO could be used for the treatment of CML-CP as it seems to push CML CD34^+^ cells into cell cycle (Figure 3e) and drives CML LSC into differentiation, as suggested by the sequencing results (Figure 5e). This might be induced by an increase in reactive oxygen species (ROS) [44]. However, GO induces DNA damage in CML cells so when the leukaemic clone is not eradicated, it might increase the risk of additional mutations that could lead to clonal evolution, treatment resistance or disease progression [25,45-47]. Thus, it might be unfavourable for patients with a good risk profile and long term follow up is required to further evaluate this.

Interestingly, PPIF is a key regulator of autophagy [48] and ROS/ischemia mediated necrosis through interaction with p53 [15]. Conversely, PPIF interacts with BCL2 to inhibit apoptosis, potentially through the inhibition of cytochrome C release [49]. Thus, PPIF is required for necrosis, but it also prevents apoptosis. High expression levels of *PPIF* might be explained by the increase in mitochondrial mass [50] and reduced necrosis-driven cell death by the reduced activity of p53 in CML LSC [13]. CsA has been used to block the interaction of PPIF with both p53 [15] and BCL2 [49], providing an anti-necrotic and pro-apoptotic effect. This is in line with our CFC results. While healthy HSC are more sensitive to CsA in the absence of IM (as they lack the pro-survival signalling from BCR-ABL1 TK), the combination of CsA and IM increases cell death in CML CFC. Although inhibition of PPIF might be a potential therapy in CML, CsA is not specific and its EC_50_ in our laboratory was not a clinically achievable concentration. Thus, more targeted compounds would need to be developed.

## 8 Conclusion

In summary, we have confirmed the existence of a TKI-independent transcriptomic programme in CML CD34^+^ cells that is not present in healthy HSC. We have also demonstrated that TKI-independent genes can be used as a novel approach for target identification in CML LSC.

## 9 Author contributions

E.G.C. performed most experiments, analysed the data, performed statistical analysis, interpreted the results, and wrote the manuscript. C.M. performed experiments. J.B.S. supported data analysis. D.V. interpreted results. M.C. interpreted results and obtained funding. S.R. and L.H. interpreted results, obtained funding, supervised the study, and performed some data analysis. H.J. and T.H. interpreted results, obtained funding, and supervised the study. All authors reviewed the manuscript.

## 10 Acknowledgements

The authors thank all those individuals with CML who generously provided samples. This study was supported by Blood Cancer UK (ref: 11017), the Howat Foundation, Glasgow Experimental Cancer Medicine Centre (funded by Cancer Research UK and the Chief Scientist’s Office, Scotland). The authors would also like to thank Pfizer for kindly providing gemtuzumab-ozogamicin for the current study. The authors would like to thank Dr Alan Hair and Ms Jennifer Cassels for the processing of primary samples, cell sorting and preparation of reagents.

## 11 Conflict of interest/disclosures

M.C. has received research funding from Cyclacel and Incyte, is/has been an advisory board member for Novartis, Incyte, Jazz Pharmaceuticals and Pfizer and has received honoraria from Astellas, Novartis, Incyte, Pfizer and Jazz Pharmaceuticals.

## Supplementary Figures

**Figure S1.**
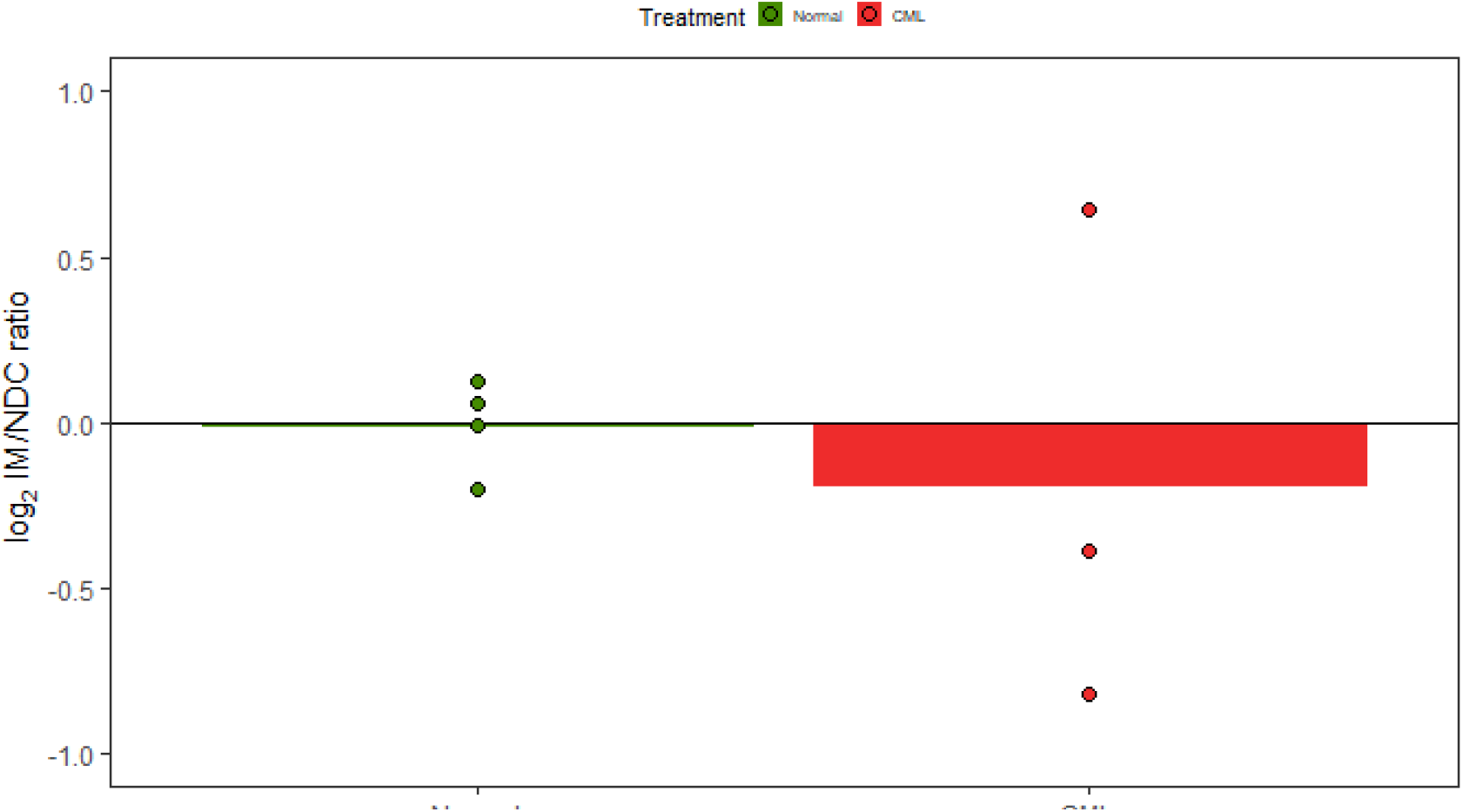
Bar plot showing CD33 expression levels on the cell surface of the cells by flow cytometry.

**Figure S2.**
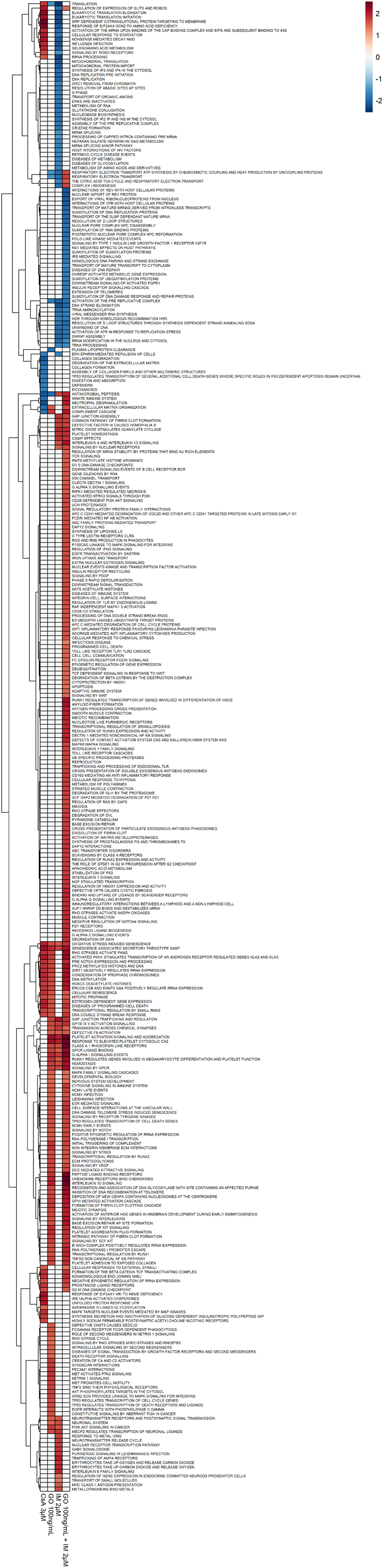
GSEA enrichment scores of all the Reactome pathways significantly enriched in at least one treatment condition. White squares represent non-significant pathways (independently of the enrichment score). Blue represents downregulated pathways while red represents upregulated pathways.

**Figure S3.**
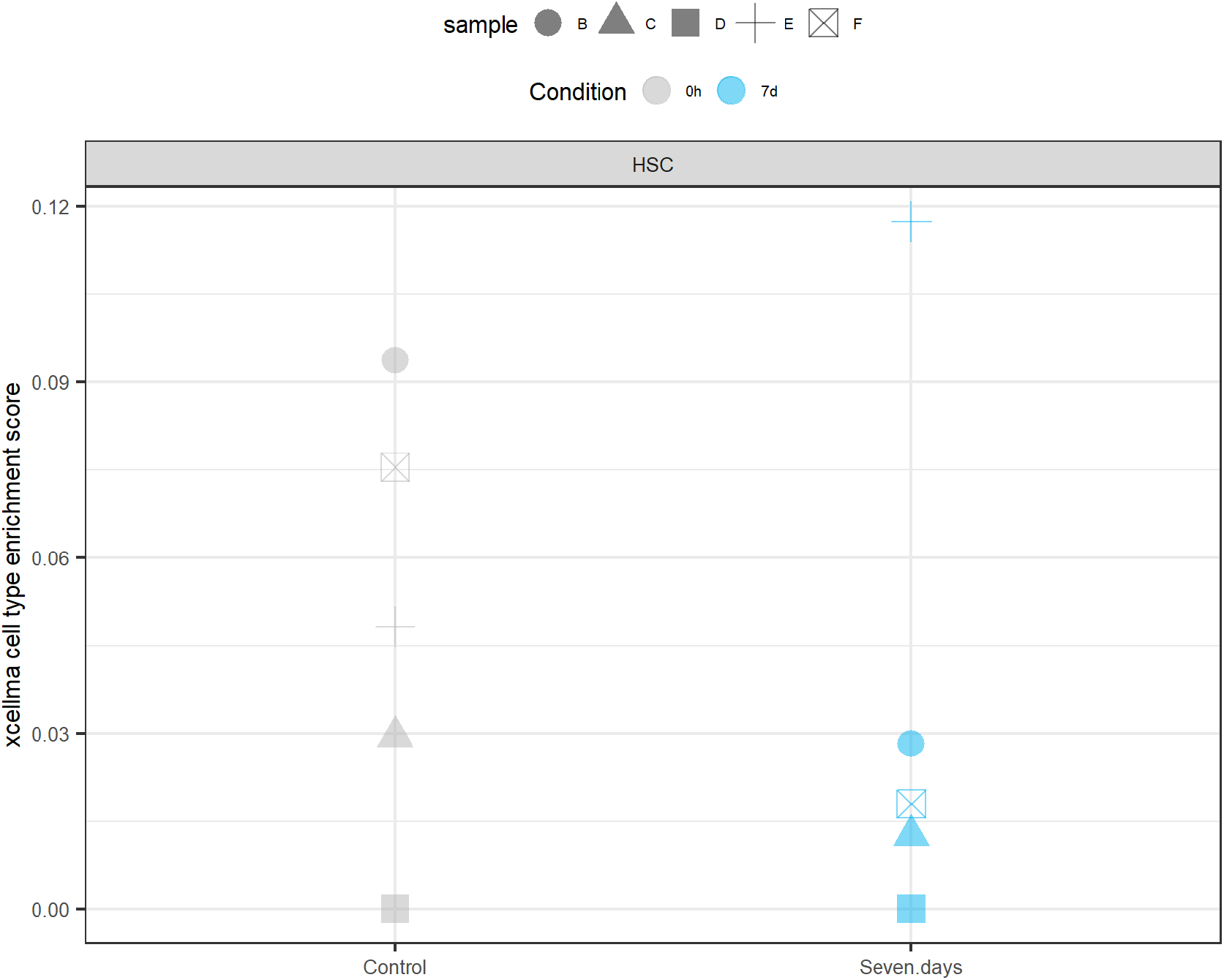
xCell enrichments for HSC in TKID. No significant difference was observed between the samples before and after treatment. Each shape represents an individual patient while each colour represents an individual time-point.

**Figure S4.**
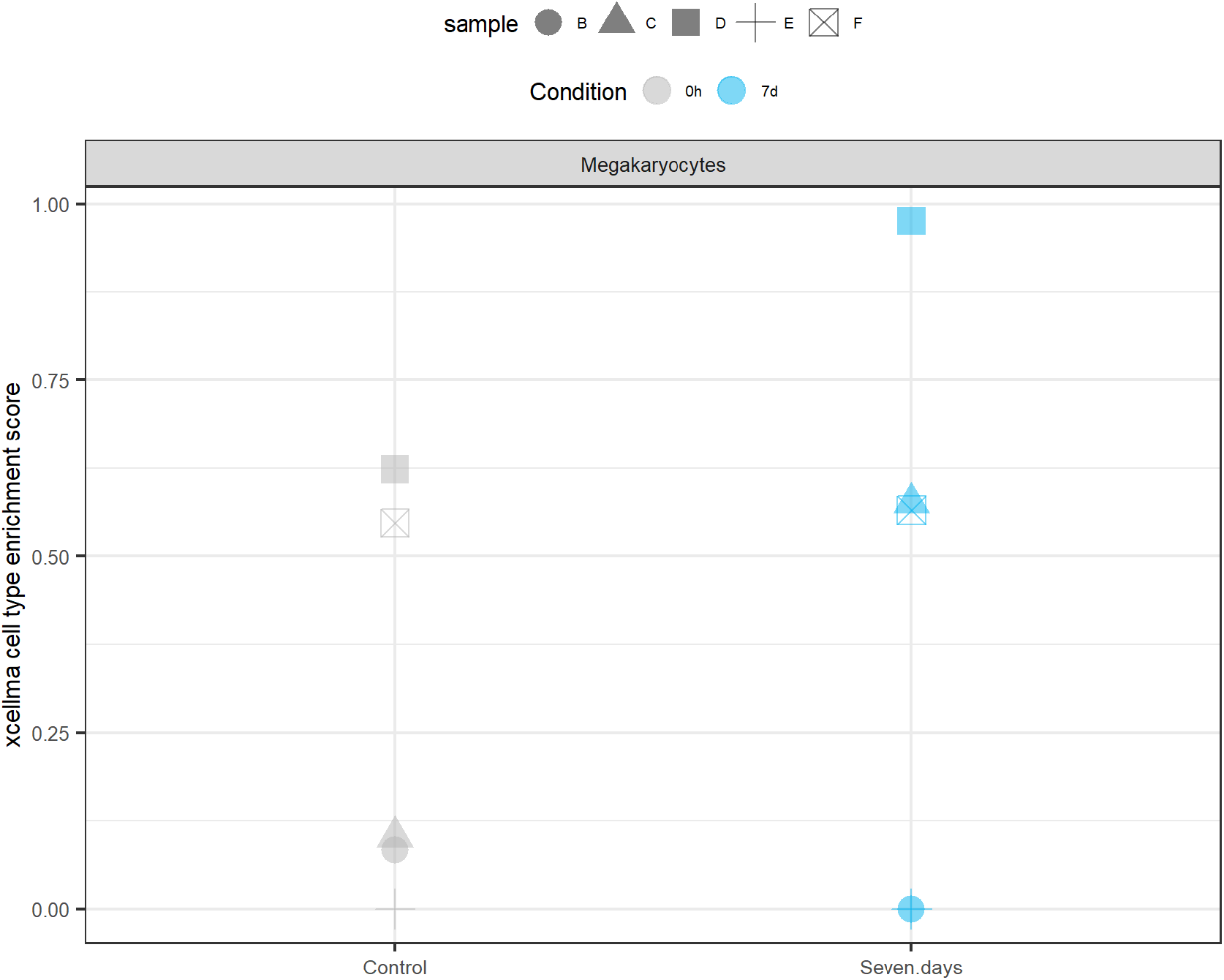
xCell enrichments for megakaryocytes in TKID. No significant difference was observed between the samples before and after treatment. Each shape represents an individual patient while each colour represents an individual time-point.

## Supplementary tables

**Table S1. Summary of the CML primary patient samples used in this project**. The levels of BCR-ABL1 by qPCR refer to the 6 months checkpoint. n/a not available.

**Table S2. Summary of the nCML samples used in this project**. n/a not available.

**Table S3**. List of TKIi genes with log_2_ fold-changes of datasets TKID and CMLD1. The tables also states if the genes were also analysed by qPCR.

**Table S4**. Summary of the number of differentially expressed genes in the RNAseq experiment for each of the treatment conditions compared to the no drug control. The table also states the number of TKIi genes differentially expressed for each condition.

**Table S5**. List of differentially expressed genes of the combination treatment 100ng/mL GO + 2μM IM compared to no drug control.

**Table S6**. List of differentially expressed genes in GO (100ng/mL) treated cells compared to no drug control.

**Table S7**. List of differentially expressed genes in CsA (3μM) treated cells compared to no drug control.

**Table S8**. List of differentially expressed genes in 2μM IM treated cells compared to no drug control.

**Table S9**. Statistics of the pairwise t-tests performed for the enrichment of HSC using xCell on the bulk RNAseq. The shown t-tests are the only ones performed for the analysis.

**Table S10**. Statistics of the pairwise t-tests performed for the enrichment of megakaryocytes using xCell on the bulk RNAseq. The shown t-tests are the only ones performed for the analysis.

## Notes

https://www.ncbi.nlm.nih.gov/geo/query/acc.cgi

